# LncRNA LINC00941 Links Oncogenic KRAS Signaling to Aggressiveness and Chemoresistance in Pancreatic Cancer

**DOI:** 10.1101/2025.09.29.679231

**Authors:** Thalita Bueno Corrêa, Gabriel Lucas da Fonseca, Rauni Borges Marques, Sandro Mascena Gomes-Filho, Lutero Hasenkamp, Yuli Magalhães, Lillian Russo, Nicolas Carlos Hoch, Fabio Luis Forti, Elena Levantini, Eduardo Moraes Reis, Daniela Sanchez Bassères

**Author notes:** **Send correspondence to:** Daniela S. Bassères, Universidade de São Paulo, Instituto de Química, Departamento de Bioquímica, Av. Prof. Lineu Prestes 748, Bloco 12 inferior, sala 1200, 05508-000, São Paulo, SP, Brazil.

## Abstract

Pancreatic ductal adenocarcinoma (PDAC) is one of the deadliest cancers, driven largely by oncogenic *KRAS*, yet effective targeted therapies remain unavailable. Long noncoding RNAs (lncRNAs) are emerging as key regulators of tumor biology, but their role in *KRAS*-driven PDAC is not well defined. To address this gap, we integrated RNA sequencing, exome, and clinical data from The Cancer Genome Atlas (TCGA) and identified 49 long intergenic noncoding RNAs (lincRNAs) differentially expressed according to *KRAS* status. Among the ten most abundant, LINC00941 and AC006262.5 showed the strongest differential expression in an independent PDAC cohort from the International Cancer Genome Consortium (ICGC). Experimental validation in *KRAS*-mutant PDAC cell lines and isogenic pancreatic epithelial models confirmed *KRAS*-dependent regulation of LINC00941, which was consistently upregulated in patient tumors and correlated with poor prognosis in both TCGA and ICGC datasets. Single-cell transcriptomic analysis further demonstrated that LINC00941 is enriched in malignant epithelial populations. Functional assays revealed that LINC00941 silencing impaired migration and invasion, reduced DNA repair capacity, and sensitized PDAC cells to gemcitabine, while having little effect on viability. Supporting these findings, co-expression and enrichment analyses linked LINC00941 to pathways regulating cell adhesion, motility, extracellular matrix organization, and DNA repair. Together, these findings demonstrate that oncogenic *KRAS* reshapes the lncRNA landscape in PDAC and identify LINC00941 as a *KRAS*-regulated oncogenic lncRNA that promotes aggressiveness and chemoresistance, highlighting its prognostic value and potential as a therapeutic target.

## Introduction

Pancreatic ductal adenocarcinoma (PDAC) is the most common malignancy of the pancreas, accounting for approximately 90% of cases and is an extremely lethal disease, with a median survival of less than 1 year and a 5-year overall survival rate of less than 15% (1–3). This bad prognosis stems from an inability to detect PDAC in early stages, its aggressive metastatic dissemination and a lack of effective targeted therapies. The most common molecular mechanism that drives PDAC growth and maintenance is constitutive activation of the KRAS GTPase by oncogenic mutations, which are widespread in PDAC, occurring in over 90% of patients (4). Even though direct KRAS inhibitors have been recently approved for KRAS^G12C^-mutant lung cancer (5), KRAS^G12C^ mutations are found in less than 2% of PDAC cases (6) and to date there are no specific *KRAS*-driven therapies approved for PDAC patients (4). In addition, PDAC tumors often display intrinsic or acquired resistance to conventional chemotherapy, as well as to KRAS inhibitors (4,6,7). In order to overcome these challenges, it is important to better understand the molecular mechanisms employed by KRAS to promote malignant transformation, metastatic spread and chemoresistance, thereby allowing for the identification of potential new KRAS targets that could be explored in combination with KRAS inhibitors, not only for the development of new anti-tumor therapeutic approaches for PDAC, but also that could serve as biomarkers for early PDAC detection and disease monitoring.

Long noncoding RNAs (lncRNAs) are defined as transcripts longer than 200 nucleotides that do not code for proteins. These transcripts can arise from various genomic regions, such that lncRNA genes can overlap protein-coding genes, reside within introns of protein-coding genes or be located in intergenic regions of the genome. In line with their genomic diversity, lncRNAs form a broad class of RNA molecules that can fold into diverse, complex secondary and tertiary structures capable of interacting with DNA, RNAs and proteins. As a result, lncRNAs play a wide array of important biological roles (8,9). In cancer, the expression of lncRNAs is often dysregulated, and aberrantly expressed lncRNAs have been linked to all major cancer hallmarks, clinical outcomes, and a subset of lncRNAs display bona-fide oncogenic or tumor suppressive functions (10–12). In addition, lncRNAs are typically expressed at lower levels than mRNAs, which makes lncRNAs promising therapeutic targets, as they could be targeted with lower drug doses, potentially reducing off-target toxicity (9). Because their expression is often highly tissue-specific, coupled to the fact that lncRNAs can be found in exossomes and can be detected in serum and urine samples, lncRNAs are strong candidates for use as cancer biomarkers (9,13). Nonetheless, despite their emerging role in cancer biology, our understanding of KRAS-regulated lncRNAs involved in promoting PDAC malignant behaviour remains incomplete.

In this study, we used a transcriptomic approach to profile lncRNA expression according to *KRAS* mutation status in PDAC and identified LINC00941 as a *KRAS*-regulated oncogenic lncRNA linked to poor patient outcome. By establishing its regulation by *KRAS* and its association with aggressive tumor features, our study positions LINC00941 as both a mechanistic contributor to PDAC progression and a promising target for therapeutic intervention.

## Material and methods

### Differential expression analysis according to KRAS mutation status in PDAC gene expression datasets

PDAC RNA-seq data from TCGA (PAAD-US, 144 cases) and ICGC (PACA-AU, 98 cases) were retrieved from the ICGC data portal (https://dcc.icgc.org), and corresponding *KRAS* mutation data for both datasets was retrieved from cBioPortal. For differential expression analysis, TCGA raw read counts were aligned to the GRCh38.p13 genome using RSEM (14) and normalized in R. DESeq2 (15) was used to compare *KRAS* mutant (KRAS^mut^, n=131) versus wild-type (KRAS^WT^, n=14) samples. Differentially expressed genes were filtered for lincRNAs with |fold change| ≥ 2 and adjusted p-value ≤ 0.05.

For downstream analyses, CPM-normalized counts were transformed using log_2_(CPM + 1) and used to compare lincRNA expression between KRAS^mut^ and KRAS^WT^ samples in both datasets. TCGA log_2_(CPM + 1) values were also used to examine expression across specific *KRAS* mutation subtypes. Statistical significance was determined using the nonparametric Wilcoxon rank-sum test, with p < 0.05 considered significant.

### Differential lncRNA expression analysis in PDAC tumors and adjacent pancreatic tumors

We have previously performed a comprehensive lncRNA differential expression analysis on a dataset comprised of 14 PDAC patient-matched tumor and non-tumor samples (14). Therefore, for this analysis, we simply retrieved the differential expression data for the lincRNAs identified in this study. A comprehensive description of the differential expression analysis performed for this dataset can be found in Paixão et al (14).

### PDAC single-cell lincRNA expression analysis

A processed count matrix with single-cell RNAseq expression data from 41,986 cells from 24 PDAC and 15,544 cells from 11 nontumor pancreatic samples generated by Peng et al (15) was retrieved from Genome Sequence Archive (accession number CRA001160). The original data was already filtered, and only cells with more than 200 detected genes and <10% mitochondrial RNA content were kept. The “Seurat” package (v. 5.0.3) in R (16) was used for normalization, selection of informative features, scaling, dimensionality reduction, and clustering of cells into different cellular identities, and annotation using the cell markers described by Peng et al (15). The normalized log-transformed count frequency in individual cells in each cluster was used to compare expression of LINC00941 and co-expressed genes among the different cell clusters. Enrichment dot plots were generated with the Seurat “DotPlot” function. In these plots, each gene is represented by two dots (ductal-1 and ductal-2). Dot size corresponds to the percentage of cells in the cluster with nonzero expression of the gene, and dot color represents the scaled average expression (“enrichment score”), ranging from –0.4 (low) to +0.4 (high). Differential expression analysis between ductal 1 and ductal 2 epithelial cells was performed using the Wilcoxon method. Genes were considered differentially expressed if they showed a log_2_FC ≥ |0.58| and an adjusted p-value ≤ 0.05.

### LINC00941 co-expression and functional enrichment analysis

RNA-seq count data from the TCGA-PAAD-US cohort were analyzed to identify genes co-expressed with LINC00941. Raw gene-level counts were imported into R and processed with edgeR (v4.4.2). Genes with missing values were excluded, and counts per million (CPM) were calculated. Lowly expressed genes were filtered out using filterByExpr function in edgeR. LINC00941 passed the filtering threshold and was retained. Filtered counts were variance-stabilized by log₂ transformation [log₂(CPM + 1)].

The expression profile of LINC00941 was extracted, and pairwise Spearman correlation coefficients were computed between LINC00941 and all remaining genes. Correlation testing was performed with the *cor.test* function in R, and p-values were adjusted for multiple comparisons using the Benjamini–Hochberg false discovery rate (FDR) method. Genes with FDR-adjusted p-values < 0.05 were considered significantly correlated with LINC00941. The top 300 genes most strongly correlated with LINC00941 (based on Spearman correlation coefficients) were selected for functional enrichment analysis. Gene set enrichment was performed using the Enrichr R package (v3.4) considering annotations from Gene Ontology (17), KEGG (18) and Reactome (19) databases. A significance threshold of adjusted p-value of *p* < 0.05 was used to select enriched terms.

### Survival analysis

PDAC cases with incomplete clinical information regarding vital status, overall survival or follow-up time were excluded from survival analysis, after which 87 cases from the ICGC-PACA-AU dataset and 141 cases from the TCGA-PAAD-US dataset remained for analysis. For each sample, raw read counts from Ensembl annotated genes (version 89) were normalized to counts per million (CPM) units. Kaplan-Meier (KM) survival curves were generated using the R survival package (20). For each gene of interest, patients were grouped into two expression groups (high and low), according to the median expression across all samples. Differences in KM survival curves of high and low expression groups were inferred by the log-rank test with a significance threshold of p < 0.05. Hazard ratios (HR) with a 95% confidence interval were estimated using Cox regression and refer to the relative risk of death in the high expression group.

### Quantitative real-time PCR (qRT-PCR)

Total RNA was isolated using Trizol reagent (Invitrogen) following the manufacturer’s protocol. RNA (1 or 0,5 μg) was used to synthesize cDNA in a reaction volume of 20 μl using Superscript III reverse transcriptase (Invitrogen), and diluted 5 times to a final volume of 100 μl. Real-time PCR analysis using 3 μl of the cDNA was performed in a GeneAmp® 5700 Sequence Detection System (Applied Biosystems®) using 6 μl of a SYBR® Green master mix (Thermo Fisher Scientific) and 400 nM of gene-specific forward and reverse primers designed using Primer3web version 4.0.0 tool (http://bioinfo.ut.ee/primer3/). The sequences used for each primer pair are listed in **Supplementary Table 1**. Relative quantitation was determined by the ΔΔCt method using GAPDH or HMBS as endogenous controls.

### Cell culture and patient samples

Cell passages were kept to a minimum and none of the cells were passaged continuously for more than six months. Human pancreatic cancer cell lines harboring *KRAS* mutations AsPC-1 (*KRAS*^G12D^; ATCC CRL-1682) and PANC1 (*KRAS*^G12D^; ATCC CRL-1469) were grown in DMEM (Invitrogen Corp) supplemented with 10% (vol/vol) fetal bovine serum (FBS). HPDE and HPDE-KR isogenic cells were grown in Keratinocyte-SFM supplemented with EGF and BPE (Thermofisher). Primary tumor samples as well as non-malignant adjacent pancreatic tissues from PDAC patients used for RNA isolation were collected with informed consent and kept in liquid nitrogen immediately after surgical resection, as part of the tumor tissue collection maintained by the AC Camargo Cancer Center. Authorization for their use in this study was granted by the Ethics Committee of the Institution (CAAE: 15059213.0.0000.5432).

### siRNA transfections

Cells were seeded in 6-well plates at a density of 1 × 10^5^ cells/ well 24 h before transfection. siRNA transfections were performed with 4 μl of the Lipofectamine 3000 transfection reagent and 50 nM of either a non-targeting siRNA control or siRNA SMART pools targeting LINC00941, or KRAS (siGENOME siRNA, Dharmacon) according to the manufacturer’s instructions. Cells were collected for extraction of RNA and protein at 48, 72 or 96 h after transfection. For migration and invasion analyses, cells were collected 96 h after transfection.

### Western blotting

Whole cell lysates of siRNA-transfected cells were prepared 72 h post-transfection using RIPA buffer (20 mM Tris–HCl pH 7.5, 150 mM NaCl, 1 mM Na2EDTA, 1 mM EGTA, 1% NP-40) containing protease and phosphatase inhibitors (1 μg/ml leupeptin, 1% sodium deoxycholate, 2.5 mM sodium pyrophosphate, 1 mM β-glycerophosphate, 1 mM Na3VO4). Aliquots of total lysate (30 μg protein/lane) were separated in 12 or 15% polyacrylamide minigels using a MiniPROTEAN® vertical electrophoresis cell (Bio-Rad). Electrophoresis was conducted with running buffer containing 25 mM Tris, 190 mM glycine and 0.1% SDS at 120 V for 60– 90 min. Next, proteins were transferred to nitrocellulose membranes (Bio-Rad) using Towbin buffer (25 mM Tris, 192 mM glycine, 20% methanol) at 350 mA for 2 h and 30 min. The blots were blocked with 5% BSA in TBST (20 mM Tris, pH 7.5, 150 mM NaCl, 0.1% Tween-20) for 1 h and incubated with primary antibodies, followed by incubation with horseradish peroxidase (HRP)-conjugated secondary antibodies. Chemiluminescence detection was performed using an a ChemiDoc MP Imaging System (Bio-Rad). The following primary antibodies were used: anti-GAPDH (Santa Cruz Biotechnology), anti-β-tubulin (Santa Cruz Biotechnology), anti-PanRAS (Merck Millipore). All primary antibodies were used at a dilution of 1:1000 in TBST, containing 5% BSA and 0.1% NaN3, and incubated for 16 h at 4 °C. The following secondary antibodies were used at a dilution of 1:3500 in TBST, and incubated for 1 h at room temperature: anti-rabbit IgG HRP (Promega) or anti-mouse IgG HRP (Promega).

### Transwell migration and invasion assays

Transwell assays were performed using uncoated (migration assay) or Matrigel-coated (invasion assay) 24-well transwell inserts with 8 μm pore size membrane filters (Corning®). AsPC-1 cells were transfected with siRNAs as described and 72 h post-transfection 1 × 10^5^ AsPC-1 cells were resuspended in 300 μl of serum-free medium and added to the upper chamber of the transwell insert. Complete medium (500 μl) was added to the lower chamber as chemoattractant and cells were incubated for 24 h at 37 °C in 5% CO 2. Non-migrating cells were removed from the upper surface of the membrane using a cotton swab, after which migrating/invading cells on the bottom surface were stained with crystal violet. Images were obtained under an IX51 Inverted Microscope (Olympus). Cells from three random fields from three independent experiments were counted using ImageJ software.

### Cell death assay

Cell death was examined using a propidium iodide (PI, Sigma-Aldrich) and Hoechst 33342 (HO, Sigma-Aldrich) double-staining method. AsPC-1 cells were transfected with siRNA as described and, 24 h post-transfection, were seeded on 96-well plates at a density of 1×10⁴ cells per well. For siRNA-only experiments, cells were analyzed after 48 h to assess the effect of LINC00941 knockdown on viability in the absence of gemcitabine. For chemoresistance experiments, gemcitabine (10, 20, or 100 µM) was added 24 h after plating, and cell death was quantified after 5 days. For PI and HO staining, cells were washed three times with PBS, incubated with HO (25 μg/ml) and PI (1 μg/ml) for 10 min at 37 °C, and examined by fluorescence microscopy. Images were acquired with a custom TissueFAXS i-Fluo (TissueGnostics) system on a Zeiss AxioObserver 7 microscope using 20× Plan-Neofluar (NA 0.5) or 40× Plan-Apochromat (NA 0.95) objectives and an ORCA Flash 4.0 v3 (Hamamatsu) camera. The percentage of dead cells was determined as the fraction of PI-positive cells among all HO-stained cells. For each condition, ≥6 × 6 adjacent fields were acquired and ≥3000 cells were analyzed in triplicate using StrataQuest (TissueGnostics). For the siRNA-only experiment (no gemcitabine), percent cell death in siCTRL versus si941 samples was compared using a paired t-test. For chemoresistance experiments, percent cell death was quantified for siCTRL and si941 samples treated with gemcitabine at 0, 10, 20, or 100 µM. To address replicate-to-replicate variability and enable different biological interpretations, data were normalized in two complementary ways: (i) to the siCTRL group at each gemcitabine dose and replicate, which isolates the siRNA effect independent of gemcitabine-induced cytotoxicity, and (ii) to the 0 µM gemcitabine condition within each siRNA group, which provides a baseline for assessing relative sensitization. Statistical testing was performed with linear mixed-effects models including siRNA and gemcitabine concentration as fixed effects and replicate as a random effect. ANOVA on the mixed model tested for main and interaction effects; residuals were examined for normality (Shapiro–Wilk) and homogeneity of variance (Levene’s test). When assumptions were violated, a nonparametric aligned rank transform (ART) ANOVA was applied. Estimated marginal means (EMMs) were derived from the mixed model, and Tukey-adjusted pairwise contrasts between si941 and siCTRL were performed at each gemcitabine dose. Fold-change values of si941 relative to siCTRL were calculated from the EMMs. In all analyses, differences were considered statistically significant at *p* < 0.05.

### DNA damage and repair assay

siRNA-treated AsPC-1 cells (6 × 10^4^) were seeded in 12-well plates in duplicate, incubated for 72 h and subjected to ionizing gamma irradiation at a dose of 10 Gy. Irradiated cells were kept in culture for recovery for 15 min, 30 min, 1 h, 2 h or 4 h and subsequently processed for alkaline comet assays. Non-irradiated control cells were processed in parallel. For the comet assays, cells were trypsinized, resuspended in 0.5% (w/v) low-melting agarose, layered on top of glass slides previously prepared with a thin layer of 1.5% (w/v) agarose, covered with coverslips to ensure homogeneous spread of cells over the slide surface and incubated overnight at 4 °C in lysis solution (2.5 M NaCl, 100 mM EDTA, 10 mM Tris pH10) for membrane disruption. Next, samples were subjected to DNA denaturation (30 min, 4 °C) and subsequent electrophoresis in alkaline pH buffer (300 mM NaOH, 1 mM EDTA, pH > 13) at 25 V and 300 mA for 30 min at 4 °C. After electrophoresis, the slides were immersed in neutralization solution (0.4 M Tris, pH 7.5) for 15 min followed by fixation in absolute ethanol for 5 min and staining with EtBr (20 μg/ml) for DNA visualization. Slide images were captured in a fluorescence microscope Olympus BX51 and analyzed using Andor Komet 6.0 software (Oxford Instruments Inc.), which measures the amount of genomic DNA damage expressed as “Olive tail moment units”, i.e., the product of the tail length and the fraction of total DNA in the tail. One hundred cells were examined in each condition/time point (50 cells/replicate).

### Statistical analysis

All values are presented as mean ± SD or as representative images from independent experiments, with the number of replicates indicated in figure legends. For pairwise comparisons, Student’s *t* test was used for normally distributed data and the Wilcoxon rank-sum test for non-normally distributed data. In gene expression analyses, comparisons of fold-change (2^-ΔΔCt) values were analyzed accordingly, with effect sizes reported as Cohen’s *d* (for *t* tests) or rank-biserial correlation (for Wilcoxon tests). For comparisons involving more than two groups, differences were assessed by one-way ANOVA followed by Bonferroni’s multiple comparison test, unless otherwise specified. Statistical significance was defined as *p* < 0.05.

## Results

### Identification of differentially expressed long intergenic noncoding RNAs (lincRNAs) according to *KRAS* mutation status

To identify differentially expressed long intergenic noncoding RNAs (lincRNAs) in PDAC based on KRAS mutation status, we analyzed publicly available RNA sequencing (RNA-seq) and exome sequencing data from TCGA (n = 145). Our analysis compared PDAC patients harboring KRAS mutations (n = 131) with those exhibiting wild-type KRAS (n = 14). To refine our analysis, we focused specifically on lincRNAs, noncoding transcripts that do not overlap annotated protein-coding regions, eliminating potential ambiguities arising from overlapping coding sequences. Differential expression was determined using a threshold of |fold change| ≥ 2 and an adjusted p-value ≤ 0.05. This approach identified 49 differentially expressed lincRNAs, including 21 upregulated and 28 downregulated in KRAS-mutant patients (**Figure 1A** and **Supplementary Table 2**). For validation, we selected the 10 most abundantly expressed lincRNAs based on their mean normalized counts across all samples (**Supplementary Table 2**). Among these, 9 were significantly upregulated in KRAS-mutant patients (*LINC01133, LINC01559, RP3-340N1.2, RP11-211G23.2, LINC00941, RP11-362F19.1/LINC02432, AC006262.5, RP5-884M6.1 e RP11-470L19.2*), while *MIR3142HG* was the only lincRNA significantly downregulated (**Figures 1A and B**).

**Fig. 1.**
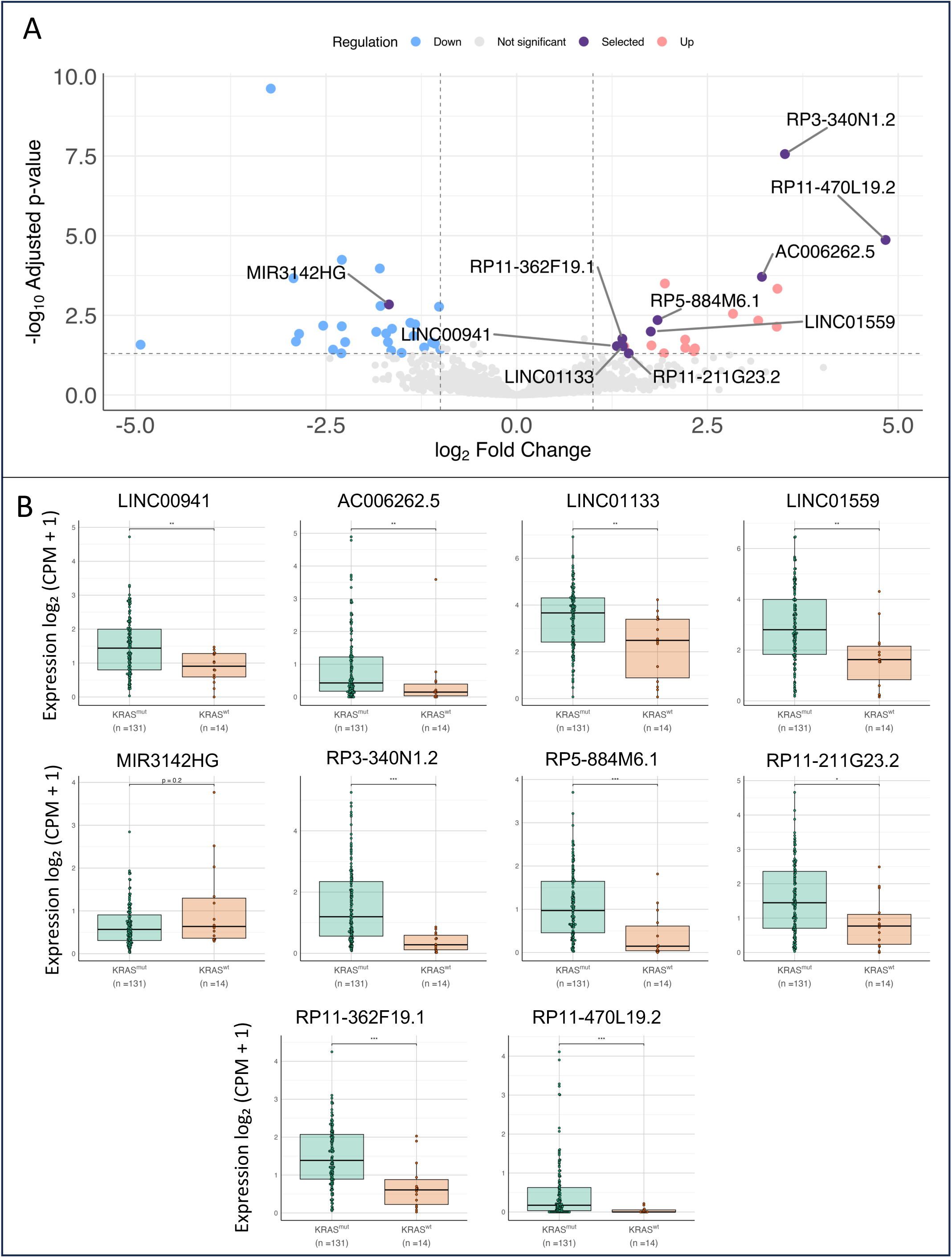
Identification of differentially expressed lincRNAs according to *KRAS* mutation status. (**A**) Volcano plot of lincRNA expression changes in the TCGA PDAC cohort. The x-axis shows the Log₂ fold change (KRAS^MUT^ vs. KRAS^WT^), and the y-axis shows the –Log₁₀ adjusted p-value. Dotted lines indicate the thresholds for significance (|Log₂FC| ≥ 1, adjusted p < 0.05). Genes are colored as significantly upregulated (up), significantly downregulated (down), not significant, or highlighted as selected abundant lincRNAs. (**B**) Dot plots comparing log₂-transformed CPM-normalized counts of the indicated lincRNAs in KRAS^MUT^ (n = 14) and KRAS^WT^ (n = 131) tumors in the TCGA PDAC cohort (PAAD-US, n = 145). Statistical significance was assessed using two-tailed Wilcoxon rank-sum tests, with p < 0.05 considered significant.

To assess whether the increased expression of the identified lincRNAs was influenced by the specific type of KRAS mutation, we reanalyzed the TCGA PAAD-US dataset (n=145), subdividing the KRAS-mutant patient group based on the specific oncogenic KRAS variants (G12D, G12R, G12V, Q61H, Q61R, and G12C). The downregulated lincRNA *MIR3142HG* only showed a significant downregulation in patients harboring KRAS Q61H mutations (n=6), indicating that the expression of this lincRNA may be influenced by the mutation type. Nonetheless, all 9 upregulated lincRNAs showed a significant upregulation in patients harboring the KRAS^G12D^ mutation (n=60) and 7 lincRNAs displayed significantly increased expression in KRAS^G12V^-mutant patients (n=38), the second most prevalent mutation in this cohort (**Figure 2**). These findings suggest that the upregulated lincRNAs are generally upregulated in KRAS-mutant samples, independent of the specific mutation type. The less common mutations did not reach statistical significance in most cases, likely due to smaller sample sizes.

**Fig. 2.**
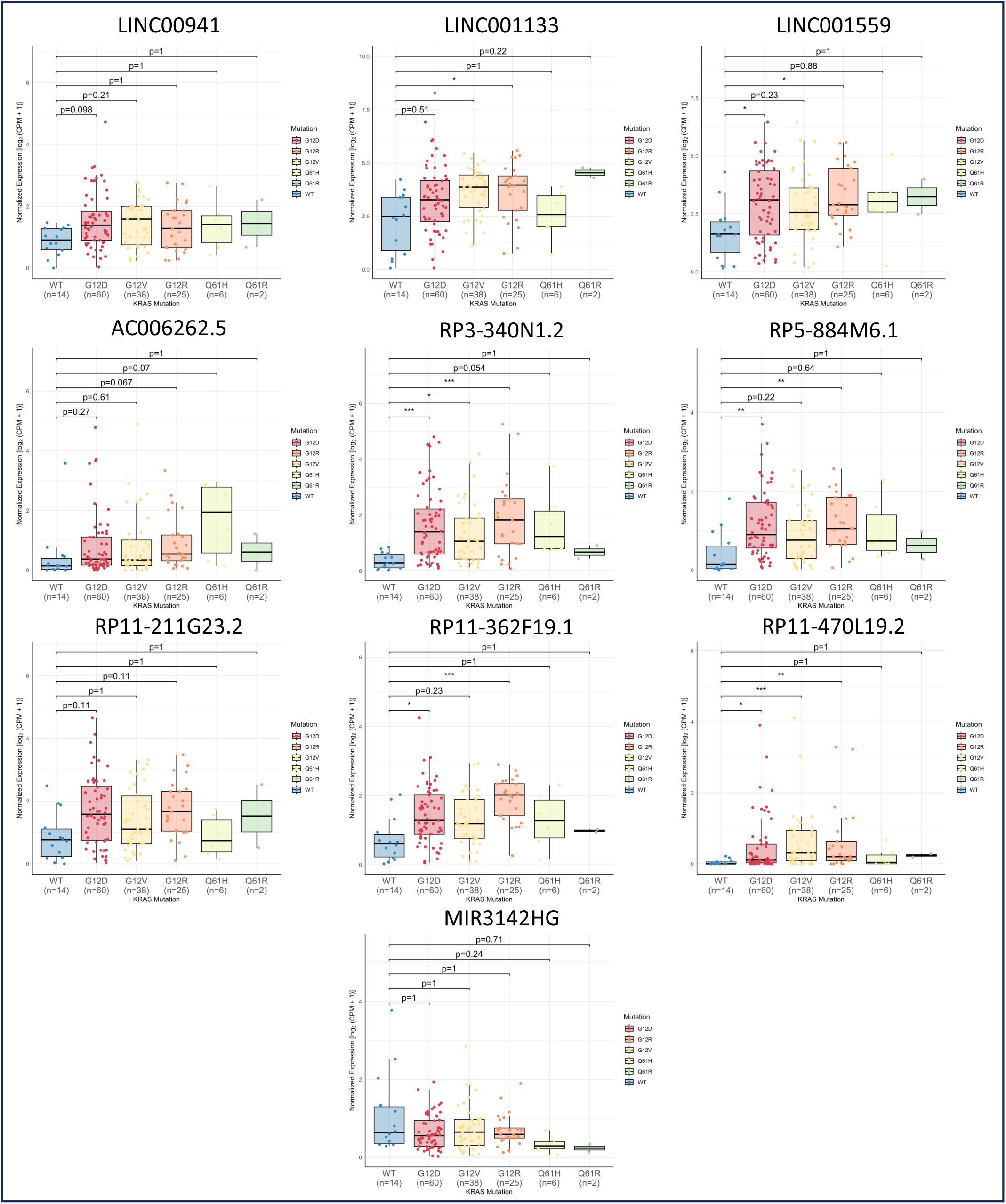
Expression of the identified lincRNAs according to *KRAS* mutation type. Dot plots showing log₂-transformed CPM-normalized counts of the indicated lincRNAs in the TCGA PDAC cohort (PAAD-US, n = 145). Expression was compared between tumors with specific KRAS mutations (G12D, n = 60; G12V, n = 38; G12R, n = 25; Q61H, n = 6; Q61R, n = 2) and wild-type KRAS tumors (KRAS^WT^, n = 14). A single case with KRAS^G12C^ (n = 1) was excluded from statistical analysis due to insufficient sample size. Statistical significance was assessed using two-tailed Wilcoxon rank-sum tests. Significance thresholds are denoted as: *p < 0.05, **p < 0.01, ***p < 0.001. p-values below the threshold are indicated.

In an attempt to validate these findings in an independent dataset, we examined the expression of these candidates in the ICGC PDAC cohort (PACA-AU, n = 98), stratifying patients by KRAS mutation status (**Supplementary Figure 2**). Although the lincRNAs showed similar expression trends in the ICGC cohort, they did not reach statistical significance, likely due to the smaller number of KRAS wild-type samples (n = 9) and the use of total RNA libraries (21), which can reduce detection sensitivity for lowly expressed lincRNAs. Among the upregulated candidates, LINC00941 (two-tailed Wilcoxon p = 0.089) and AC006262.5 (p = 0.138) showed the strongest trends toward higher expression in KRAS-mutant tumors. Based on this cross-cohort consistency in effect direction and magnitude, LINC00941 and AC006262.5 were selected for further analyses.

We started by investigating whether LINC00941 and AC006262.5 correlated with the malignant transformed phenotype of PDAC tumors. To do so, we analyzed RNA-seq data previously generated in our laboratory, which included paired tumor and adjacent non-tumor tissue samples from 14 PDAC patients (14). Both LINC00941 and AC006262.5 exhibited significantly higher expression in tumor samples compared to adjacent non-tumor tissue (**Figure 3A**). This suggests that these two lincRNAs may play a role in promoting PDAC malignancy.

**Fig. 3.**
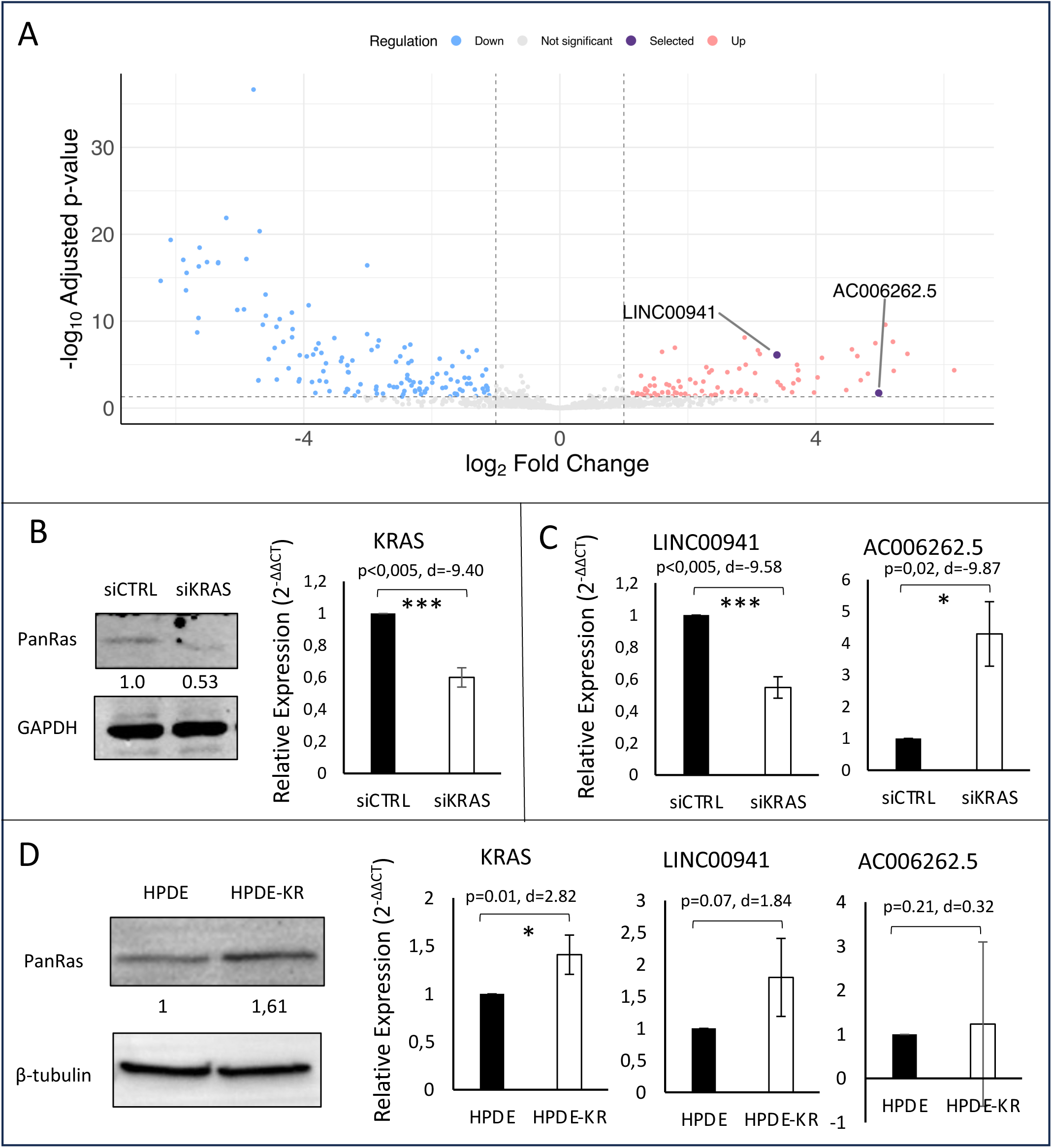
KRAS regulates the expression of lincRNAs in PDAC cell lines. **(A)** Volcano plot of lincRNA expression changes in paired PDAC tumors and adjacent pancreatic tissue (APT, n = 14). The x-axis represents log₂ fold change (PDAC vs APT), and the y-axis represents –log₁₀ adjusted p-value. Dotted lines indicate the thresholds for significance (|log₂FC| ≥ 0.58, adjusted p < 0.05). Genes are colored by significance status (upregulated, downregulated, not significant), with LINC00941 and AC006262.5 highlighted. (**B**) Validation of KRAS knockdown in AsPC-1 cells by RNA interference. *Left:* Western blot comparing Ras protein levels in siCtrl and siKRAS groups (representative of n = 3). GAPDH served as loading control. *Right:* qRT-PCR analysis of KRAS mRNA expression in siKRAS and siCtrl groups (n = 3). (**C**) KRAS-dependent regulation of candidate lincRNAs in AsPC-1 cells. qRT-PCR analysis of lincRNA expression in siKRAS and siCtrl groups (n = 3). (**D**) Expression of KRAS and lincRNAs in isogenic pancreatic cells. *Left:* Western blot comparing Ras protein levels in HPDE and HPDE-KR cells (representative of n = 1). α-Tubulin served as loading control. *Right:* qRT-PCR analysis of KRAS mRNA expression in HPDE and HPDE-KR cells (n = 3). HMBS was used as endogenous control for all qRT-PCR experiments. Statistical significance for qRT-PCR data was assessed using paired one-tailed Student’s *t*-test for KRAS knockdown validation and paired two-tailed Student’s *t*-test for target lincRNA regulation. Significance levels are denoted as *p < 0.05, ***p < 0.005. Exact *p*-values and Cohen’s *d* effect sizes are indicated.

### LINC00941 is a KRAS target in AsPC-1 cells

To investigate whether KRAS regulates LINC00941 and AC006262.5 expression in PDAC, we used siRNA-mediated knockdown of KRAS in AsPC-1 cells. Compared to control cells transfected with a non-targeting siRNA (siCtrl), cells transfected with a KRAS-specific siRNA SMARTpool (siKRAS) exhibited a ∼50% reduction in total RAS protein levels (**Figure 3B**). Given that the antibody used also detects HRAS and NRAS, the reduction in KRAS protein is likely even greater. This was corroborated by qPCR, which revealed a ∼50% decrease in KRAS mRNA levels (**Figure 3B**), confirming effective knockdown. As expected, KRAS knockdown significantly reduced LINC00941 expression by 46% (**Figure 3C**). However, contrary to our hypothesis, AC006262.5 expression increased significantly following KRAS suppression (**Figure 3C**).

To assess whether these observations were influenced by the genetic background of AsPC-1 cells, we analyzed expression of the lincRNA candidates in isogenic low-passage immortalized human pancreatic ductal epithelial cells (HPDE) and their KRAS-transformed counterparts (HPDE-KR). Despite a 41% increase in KRAS expression in HPDE-KR cells, AC006262.5 expression in HPDE-KR cells was highly variable and did not differ significantly from HPDE controls. In contrast, LINC00941 expression increased by ∼80% in HPDE-KR cells compared to controls (p = 0.07, Cohen’s d = 1.84), indicating a large effect size despite not reaching conventional statistical significance. (**Figure 3D**).

Collectively, KRAS perturbation in PDAC models affected LINC00941 expression in a manner consistent with regulation by KRAS. The effect was statistically significant in AsPC-1 cells and showed a strong trend in HPDE-KR cells (p = 0.07), supporting its role as a KRAS-regulated lincRNA.

### LINC00941 is overexpressed in PDAC tumors and in malignant PDAC cells and is associated with worse PDAC prognosis

To further investigate the association between LINC00941 and the malignant phenotype, we analyzed its expression in seven pancreatic ductal adenocarcinoma (PDAC) patient samples and five non-tumor pancreatic tissue samples by qPCR. Comparison of gene expression between non-tumor and tumor samples using the Wilcoxon rank-sum test revealed a trend toward differential expression (p = 0.073). Despite the lack of formal significance, the effect sizes were large (rank-biserial = 0.54), indicating that the magnitude of the difference between groups is biologically meaningful (**Figure 4A**). These results suggest that, although the small sample size limits statistical power, the observed expression changes may reflect genuine alterations associated with the tumor phenotype, which is consistent with our RNA sequencing data (**Figure 3A**).

**Fig. 4.**
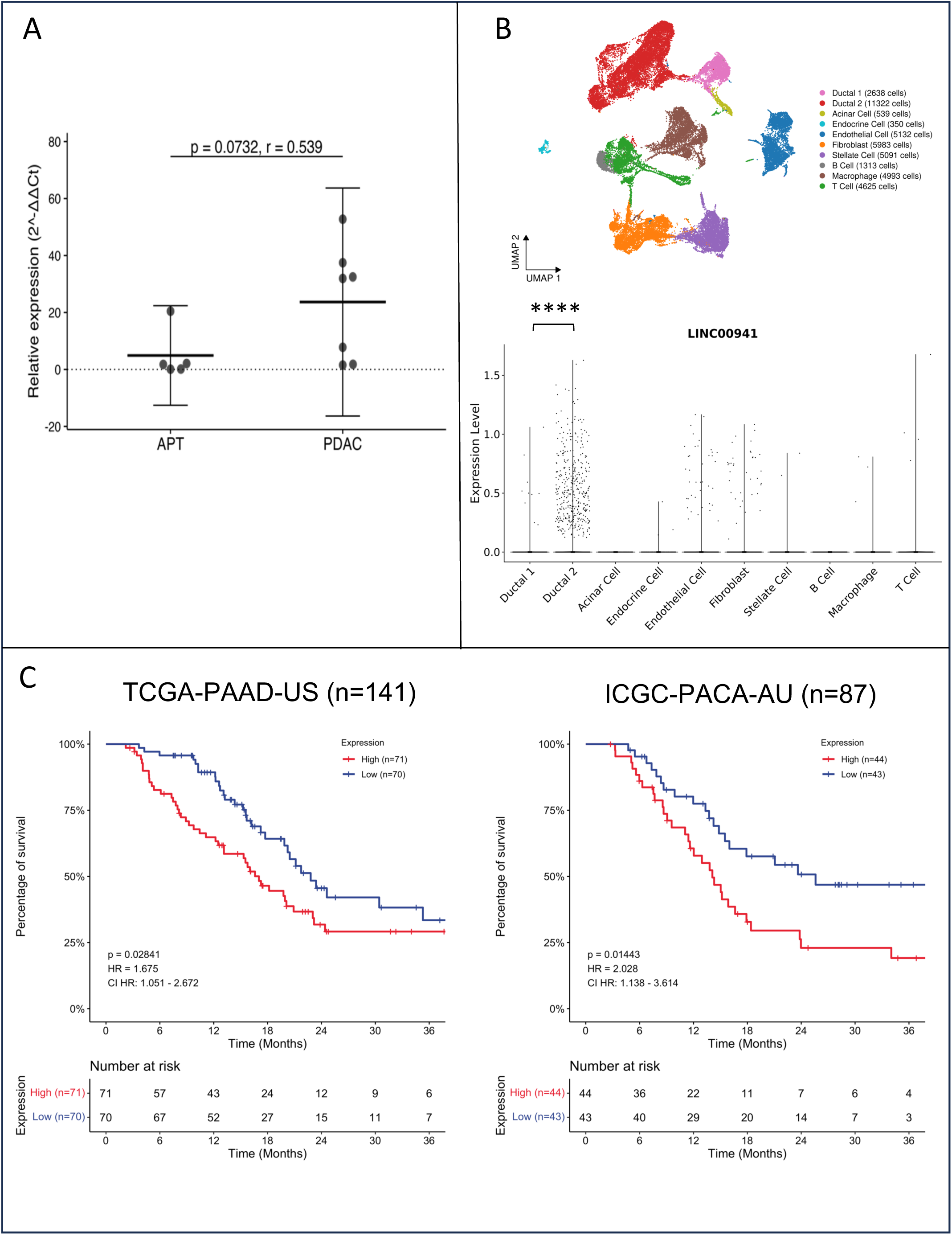
LINC00941 is overexpressed in PDAC tissue. (**A**) qRT-PCR analysis of LINC00941 expression in PDAC tumors (n = 7) compared with adjacent pancreatic tissue (APT, n = 5). Results are shown as dot plots of relative expression values, with horizontal bars indicating group means. HMBS was used as endogenous control. Statistical significance was assessed using the Wilcoxon rank-sum test; exact *p*-values and rank-biserial effect sizes are indicated. (**B**) Single-cell RNA-seq analysis of LINC00941 expression in PDAC. *Upper:* Uniform manifold approximation and projection (UMAP) plot of annotated cell clusters identified with Seurat. The number of cells per cluster is indicated. *Lower:* Violin plot showing normalized log-transformed expression level of LINC00941 across individual clusters. Differential expression testing was performed on normalized, log₂-transformed counts using the Wilcoxon rank-sum test with p-values adjusted for multiple testing (Benjamini–Hochberg), and significance level denoted as ****padj < 0.0001. (**C**) Kaplan–Meier survival analysis based on LINC00941 expression in the TCGA (PAAD-US, n = 145) and ICGC (PACA-AU, n = 87) cohorts. Patients were stratified into high- and low-expression groups using the median expression cutoff. Differences in survival were evaluated using the log-rank test; log-rank *p*-values and hazard ratios (HR) with 95% confidence intervals are shown.

Although LINC00941 was identified in bulk tumor samples, which contain not only cancer cells but also other cell types, such as stromal and immune cells, it is essential to confirm its enrichment in the malignant cell population. This is particularly relevant for PDAC, given its highly infiltrative nature, where tumors often include a substantial proportion of stromal components and normal pancreatic tissue (22).

To determine whether LINC00941 is specifically expressed in malignant PDAC cells, we analyzed single-cell RNA sequencing data (GSA, accession number CRA001160) from 24 PDAC tumor samples and 11 control pancreatic tissue samples, pre-clustered into 10 distinct cell types (15). While LINC00941 expression was detected in endothelial cells and fibroblasts, it was significantly enriched in ductal type 2 cells in contrast to ductal type 1 cells (**Figure 4B**). Notably, whereas ductal type 1 cells were found both in control pancreatic samples and in PDAC tumor samples and found to correspond to normal ductal cells, ductal type 2 cells were exclusively present in tumor samples and identified by Peng et al. (15) as the malignant PDAC cell population. These findings confirm that LINC00941 is preferentially expressed in PDAC malignant tumor cells, further supporting its role in tumor progression.

More importantly high LINC00941 expression correlates with reduced overall survival in PDAC patients from both the TCGA and ICGC cohorts (**Figure 4C**). These findings reinforce LINC00941 as the most promising KRAS-associated lincRNA candidate likely to be involved in promoting malignant tumor behavior.

### LINC00941 depletion reduces migration and invasion of KRAS-mutant PDAC cells

To investigate whether LINC00941 contributes to the KRAS-driven malignant phenotype, we silenced LINC00941 in AsPC-1 cells using siRNA and assessed its impact on cell viability, migration and invasion (**Figure 5**). LINC00941 knockdown was validated, revealing a 63% reduction in LINC00941 expression (**Figure 5A**). LINC00941 knockdown did not increase cell death (**Figure 5B**). Nonetheless, LINC00941 depletion decreased cell migration and invasion by 21% and 38%, respectively (**Figures 5C, 5D**). These findings suggest that LINC00941 enhances tumor progression by promoting a more aggressive phenotype.

**Fig. 5.**
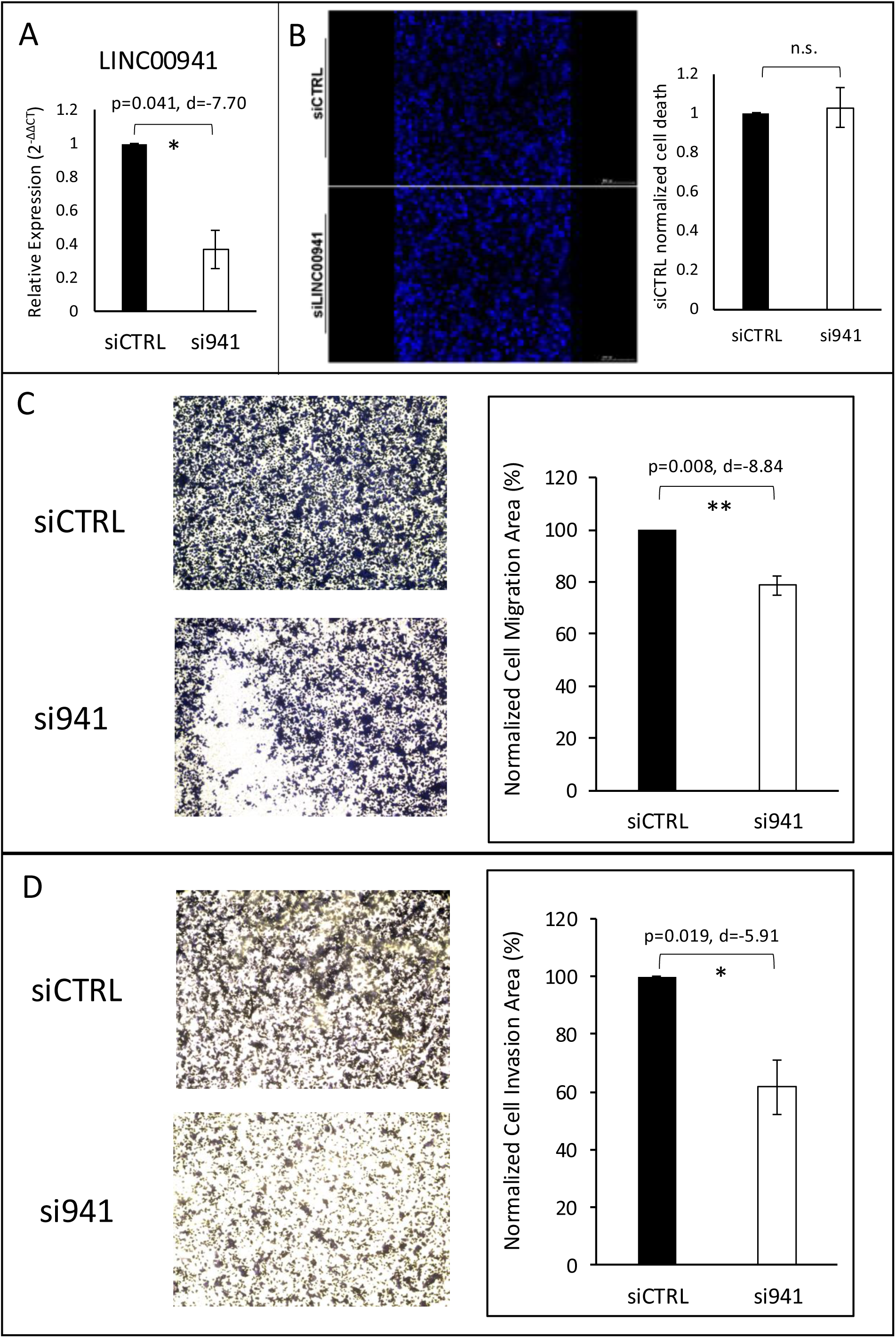
Targeting LINC00941 reduces migration and invasion of KRAS-mutant AsPC-1 cells. AsPC-1 cells were transfected with a non-targeting control siRNA (siCTRL) or a SMARTpool siRNA targeting LINC00941 (si941). (**A**) qRT-PCR analysis of LINC00941 expression in siCTRL and si941 groups (n=2). Data represent fold-change relative to siCTRL. HMBS was used as endogenous control. (**B**) Cell viability analysis. *Left:* Representative images of Hoechst- and PI-stained cells (n = 3). *Right:* Quantification of normalized cell death relative to siCTRL (n = 3). (**C–D**) Transwell migration (**C**) and invasion (**D**) assays. *Left:* Representative images (n=3). *Right:* Quantification shown as mean ± SD. Statistical significance was assessed by paired Student’s *t*-tests (one-tailed for panel A; two-tailed for panels B–D). Significance thresholds: *p* < 0.05 (*), *p < 0.01 (**),* n.s., not significant. For significant comparisons, exact *p*-values and Cohen’s *d* effect sizes are reported. Comparisons are indicated by horizontal bars.

### LINC00941 is required for efficient DNA repair

Long non-coding RNAs (lncRNAs) are frequently functionally related to protein coding genes with which they are co-expressed (23,24). Using Weighted Gene Co-expression Network Analysis (WGCNA) of RNA-Seq data from PDAC tumors and adjacent normal tissues, we previously identified LINC00941 to be co-expressed with protein coding genes associated with DNA repair (14).

To validate this association, we examined the expression of DNA repair-related genes with high connectivity to the DNA repair module of co-expressed transcripts that included LINC00941 (14), including both genes positively correlated with LINC00941 (SPIDR, PPP4C, BACH1, and ATRX), as well as negatively correlated genes (TERF21P and CDK7). We also included TOP2A, a DNA repair gene found to be positively correlated with LINC00941 in a different WGCNA module of the same data. LINC00941 knockdown significantly downregulated PPP4C, BACH1 and TOP2A. A decreasing trend was also observed for ATRX (p = 0.07, Cohen’s d = –2.28) and SPIDR (p = 0.18, Cohen’s d = –2.27), whereas TERF21P (p = 0.53) and CDK7 (p = 0.42) expression remained unchanged (**Figure 6A**). These results show that LINC00941 only affects the expression of positively correlated genes in this pathway, thus suggesting that LINC00941 does not seem to act to repress gene expression, but rather acts as a positive regulator of gene expression. The LINC00941-mediated regulation of PPP4C and BACH1 was confirmed in PANC-1 cells (**Supplementary Figure 2**). Interestingly, analysis of single cell sequencing data of PDAC tumors revealed that, of the DNA repair genes positively correlated with LINC00941, only the qPCR-validated genes (BACH, PPP4C and TOP2A) are enriched in malignant type 2 ductal cells, with ATRX and SPIDR being enriched in type 1 ductal cells (**Figure 6B**).

**Fig. 6.**
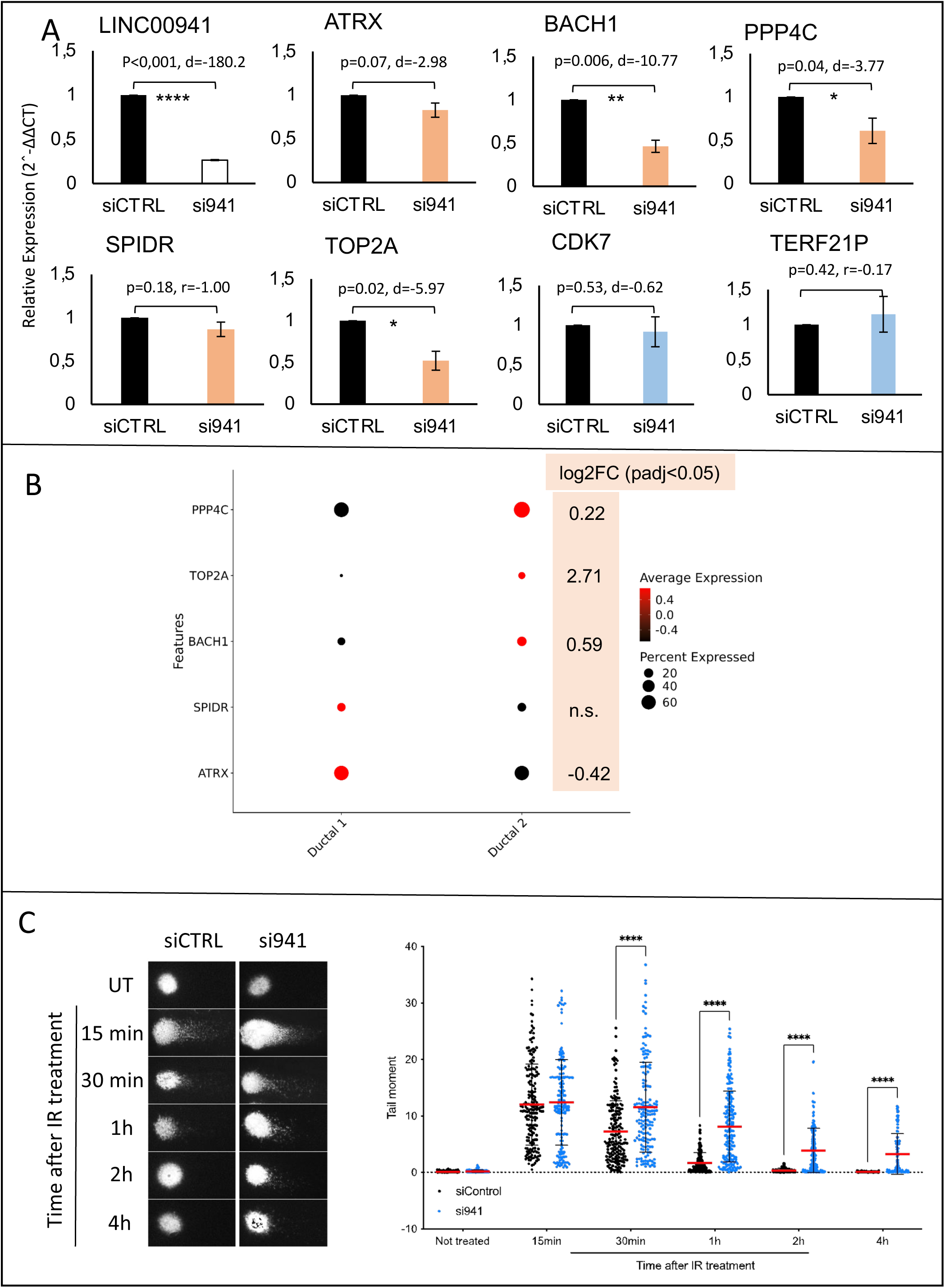
LINC00941 depletion impairs DNA repair. AsPC-1 cells were transfected with non-targeting control siRNA (siCTRL) or a SMARTpool siRNA targeting LINC00941 (si941). **(A)** qRT-PCR analysis of LINC00941 and selected DNA-repair genes 48 h after transfection (n = 3 independent experiments). Data are shown as fold-change relative to siCTRL (mean ± SD). HMBS was used as endogenous control. Bars colored red indicate genes positively correlated with LINC00941; blue indicates negatively correlated genes. Statistical testing: paired one-tailed (LINC00941) or two-tailed (DNA repair genes) Student’s *t*-test for normally distributed data (Shapiro–Wilk) and two-tailed Wilcoxon signed-rank test for non-normal data. Significance: *p* < 0.05; \*\**p* < 0.005; \*\*\**p* < 0.001. Exact *p*-values and effect sizes are reported (Cohen’s *d* for parametric tests; rank-biserial correlation for nonparametric tests). **(B)** Single-cell RNA-seq enrichment dot plot for LINC00941-correlated DNA-repair genes comparing ductal 1 and ductal 2 epithelial subpopulations. Dot size indicates the percentage of cells in that subpopulation with detectable expression as indicated; dot color indicates the enrichment score as indicated. Differential expression testing was performed on normalized, log₂-transformed counts using the Wilcoxon rank-sum test and p-values were adjusted for multiple testing (Benjamini–Hochberg), with padj < 0.05 considered significant. Log_2_FC values are indicated for significant enrichments. n.s.) not significant **(C)** Alkaline comet assay measuring DNA fragmentation. At 48 h post-transfection, cells were either left untreated (UT) or exposed to 10 Gy γ-irradiation and collected at 15, 30, 60 (1 h), 120 (2 h) and 240 (4 h) min after irradiation. *Left:* representative single-nucleus comet images (n=2). *Right:* dot plots of olive tail moment (OTM) for individual nuclei at each time point; horizontal bars indicate mean ± SD. Data derive from 50 nuclei per condition per experiment, pooled from two independent experiments (total n = 100 nuclei per condition per time point). Statistical analysis was performed using two-way ANOVA with Tukey’s post hoc test. Significance between groups is indicated by horizontal bars (\*\*\*\**p* < 0.0001).

Since DNA repair is essential for maintaining genomic integrity and promoting tumor progression, we next assessed the effect of LINC00941 knockdown on DNA damage repair following γ-radiation-induced DNA damage. While both control and LINC00941-depleted cells exhibited similar initial DNA fragmentation 15 minutes after γ-radiation exposure, LINC00941-silenced cells showed significantly delayed DNA repair at all subsequent time points indicated by higher DNA fragmentation than control cells (**Figure 6C**). Notably, at the final time point (4 hours), DNA damage in control cells had returned to baseline, whereas LINC00941-depleted cells still retained substantial DNA fragmentation. These findings confirm that LINC00941 is crucial for efficient DNA repair.

### LINC00941 contributes to gemcitabine chemoresistance

Since enhanced DNA repair is often associated with chemoresistance (25), we hypothesized that LINC00941 promotes resistance to gemcitabine, a nucleoside analog routinely used in PDAC therapy, both for advanced PDAC, as well as an adjuvant therapy for patients eligible for surgical resection. To test this, we silenced LINC00941 in AsPC-1 cells and assessed cell death following treatment with 10, 20, and 100 µM gemcitabine for 5 days (**Figure 7A**). These concentrations were chosen, because prior pharmacokinetic studies in pancreatic cancer patients using fixed-dose rate infusions have reported plasma gemcitabine concentrations of ∼15 µM. Therefore, our *in vitro* doses of 10-20 µM are within the clinically relevant range (26), with 100 µM representing a suprapharmacological dose.

**Fig.7.**
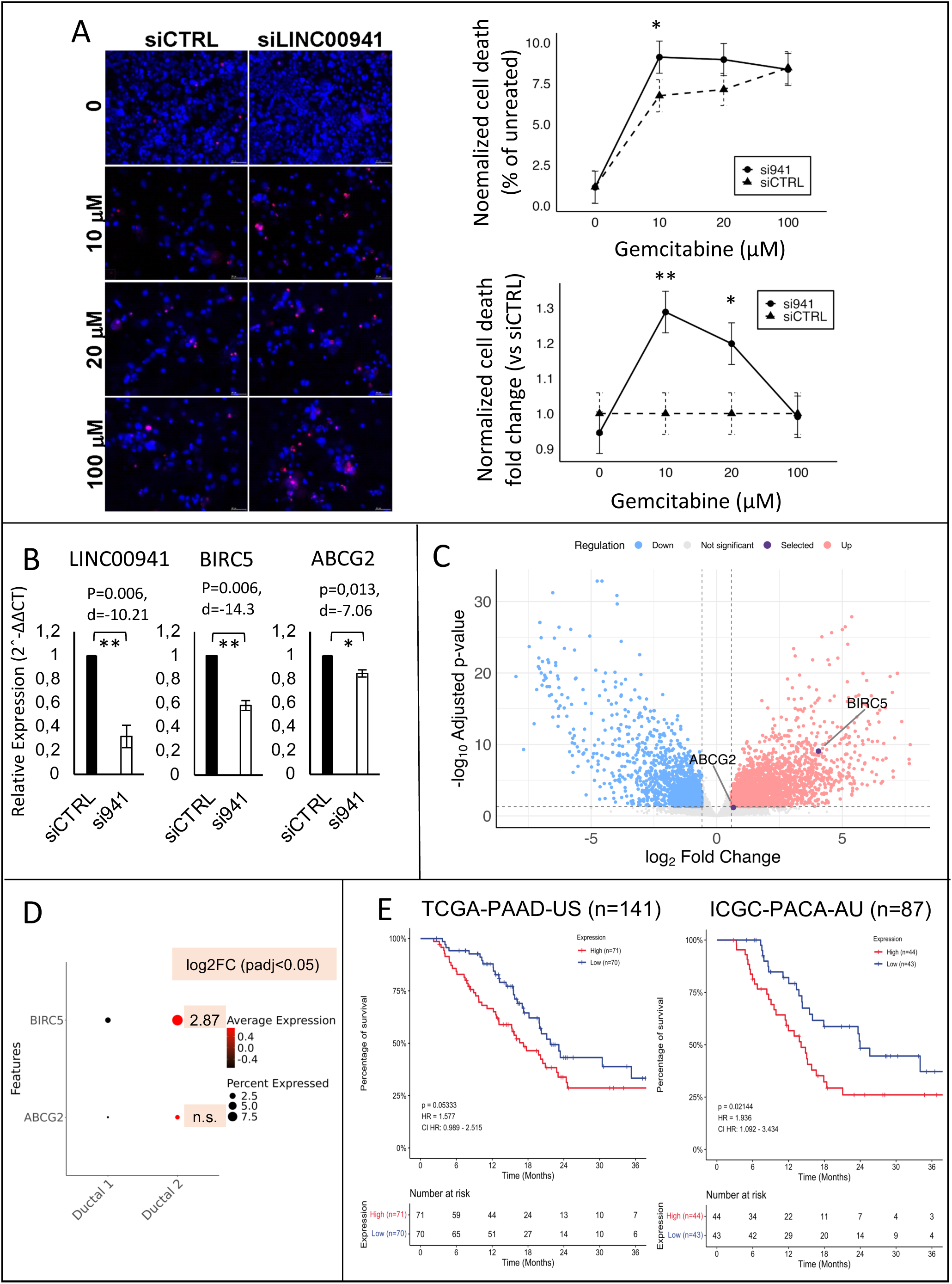
LINC00941 depletion sensitizes AsPC-1 cells to gemcitabine. AsPC-1 cells were transfected with a non-targeting control siRNA (siCtrl) or with a siRNA SMARTpool targeting LINC00941 (siLINC00941) as described in methods. After 48h, cells were treated with the indicated Gemcitabine doses for 5 days and cell death was analyzed by Hoechst/PI staining. (**A**) *Left:* Representative fluorescence images (Hoechst, blue; PI, red) from three independent experiments. *Right:* Normalized cell death graphs as mean ± SE (n = 3). *Upper:* graph represents percent dead values normalized to the 0 µM dose within each siRNA to visualize the overall gemcitabine dose-response. *Lower:* graph represents percent dead cell values normalized to the siCTRL group at each gemcitabine concentration to highlight siRNA effects independently of gemcitabine dose. Two-way aligned rank transform (ART) ANOVA, a nonparametric factorial test, detected significant main effects of siRNA (F₁,₁₄ = 10.45, p = 0.006), gemcitabine dose (F₃,₁₄ = 6.73, p = 0.005), and a significant siRNA × dose interaction (F₃,₁₄ = 7.25, p = 0.004). Post-hoc contrasts (emmeans, Tukey-adjusted) were used for pairwise comparisons in both graphs (*p<0.05, **p<0.01). (**B**) qRT-PCR analysis of LINC00941 and selected chemoresistance-related genes 48 h after transfection (n = 3). Data are shown as fold change relative to siCTRL (mean ± SD). HMBS was used as endogenous control for LINC00941 and BIRC5 and GAPDH was used as endogenous control for ABCG2. Statistical significance was assessed using paired one-tailed (LINC00941) or two-tailed (ABCG2 and BIRC5) Student’s *t*-test. Significance levels are denoted as *p < 0.05, **p < 0.01. Exact *p*-values and Cohen’s *d* effect sizes are indicated. (**C**) Volcano plot of protein coding genes expression changes in paired PDAC tumors and adjacent pancreatic tissue (APT, n = 14). The x-axis represents log₂ fold change (PDAC vs APT), and the y-axis represents –log₁₀ adjusted p-value. Dotted lines indicate the thresholds for significance (|log₂FC| ≥ 0.58, adjusted p < 0.05). Genes are colored by significance status (upregulated, downregulated, not significant), with ABCG2 and BIRC5 highlighted. (**D**) Single-cell RNA-seq enrichment dot plot for ABCG2 and BIRC5 comparing ductal 1 and ductal 2 epithelial subpopulations. Dot size indicates the percentage of cells in that subpopulation with detectable expression and dot color indicates the enrichment score. Differential expression testing was performed on normalized, log₂-transformed counts using the Wilcoxon rank-sum test and p-values were adjusted for multiple testing (Benjamini–Hochberg), with padj < 0.05 considered significant. Log_2_FC values are indicated. n.s.) not significant (**E**) Kaplan–Meier survival analysis based on BIRC5 expression in the TCGA (PAAD-US, n = 145) and ICGC (PACA-AU, n = 87) cohorts. Patients were stratified into high- and low-expression groups using the median expression cutoff. Differences in survival were evaluated using the log-rank test; log-rank *p*-values and hazard ratios (HR) with 95% confidence intervals are shown.

When cell death was normalized to the 0 µM dose, LINC00941 depletion significantly increased cell death at 10 µM gemcitabine 27.5% (p = 0.047), with a similar but non-significant trend at 20 µM (p = 0.12) (**Figure 7A, upper graph**). Importantly, when the data were normalized to the siCTRL condition at each dose, the sensitizing effect of LINC00941 knockdown was even more evident: cell death was significantly higher at both 10 µM (p = 0.0039) and 20 µM (p = 0.032) (**Figure 7A, lower graph**). At 100 µM, however, gemcitabine-induced cytotoxicity dominated, masking additional contributions from LINC00941. These results demonstrate that LINC00941 depletion sensitizes PDAC cells to gemcitabine, with the strongest effect observed at clinically relevant doses, thereby supporting a role for this lncRNA in promoting chemoresistance.

Additionally, LINC00941 knockdown reduced expression of chemoresistance-related genes ABCG2 and BIRC5 by 15% and 42%, respectively (**Figure 7B**). ABCG2 is a key drug efflux transporter implicated in PDAC chemoresistance (27) and BIRC5, also called Survivin, is a member of the Inhibitor of Apoptosis (IAP) family also found to promote PDAC resistance to gemcitabine (28). In tumor versus adjacent pancreatic tissue, both genes exhibit log_2_FC values above the differential expression threshold (log_2_FC>0.58), although only BIRC5 reached statistical significance (padj = 2.89E-34), showing a marked fold change increase (**Figure 7C**). By contrast, ABCG2 shows a similar trend (padj = 0.065) but does not meet statistical significance. Both are enriched in type 2 ductal cells in single cell analysis, with BIRC 5 displaying a significant fold change increase in type 2 ductal cells (log_2_FC= 2,87, padj= 2,89E^-34^, **Figure 7D**). Finally, high BIRC5 expression correlates with reduced overall survival in PDAC patients from both the TCGA and ICGC cohorts (**Figure 7E**). These results indicate that LINC00941 contributes to PDAC chemoresistance, potentially by enhancing DNA repair, promoting drug efflux mechanisms and inhibiting chemotherapy-induced apoptosis.

Finally, to gain further insight into the role of LINC00941 in KRAS-mutant PDAC, we performed a co-expression analysis using the TCGA cohort. For downstream analyses, we applied a cutoff of Spearman correlation coefficient ≥ 0.65, yielding 393 highly co-expressed genes with LINC00941 (**Supplementary Table 3**). Among these, 13 were also differentially expressed in KRAS-mutant PDAC patients (**Figure 8A**). Consistent with the importance of LINC00941 in KRAS-driven PDAC, single-cell analysis showed that most of these LINC00941-correlated DEGs were enriched in type 2 ductal cells compared with type 1 ductal cells (**Figure 8A**).

Functional enrichment of the top 300 LINC00941-correlated genes revealed significant associations with DNA repair, cell motility, extracellular matrix regulation, and cell adhesion pathways (**Supplementary Table 4**, **Figure 8B**). Representative genes from these categories were not only enriched in type 2 ductal cells (**Figure 8C**) but also significantly upregulated in PDAC tumors (15 out of 18, **Figure 8D**). Importantly, ITGA3 and RAD51AP1, were also significantly associated with patient survival (**Supplementary Figure 3**). Together, these findings reinforce our experimental results and support a model in which LINC00941 promotes tumor aggressiveness and contributes to chemoresistance and poor prognosis in KRAS-driven PDAC.

**Fig. 8.**
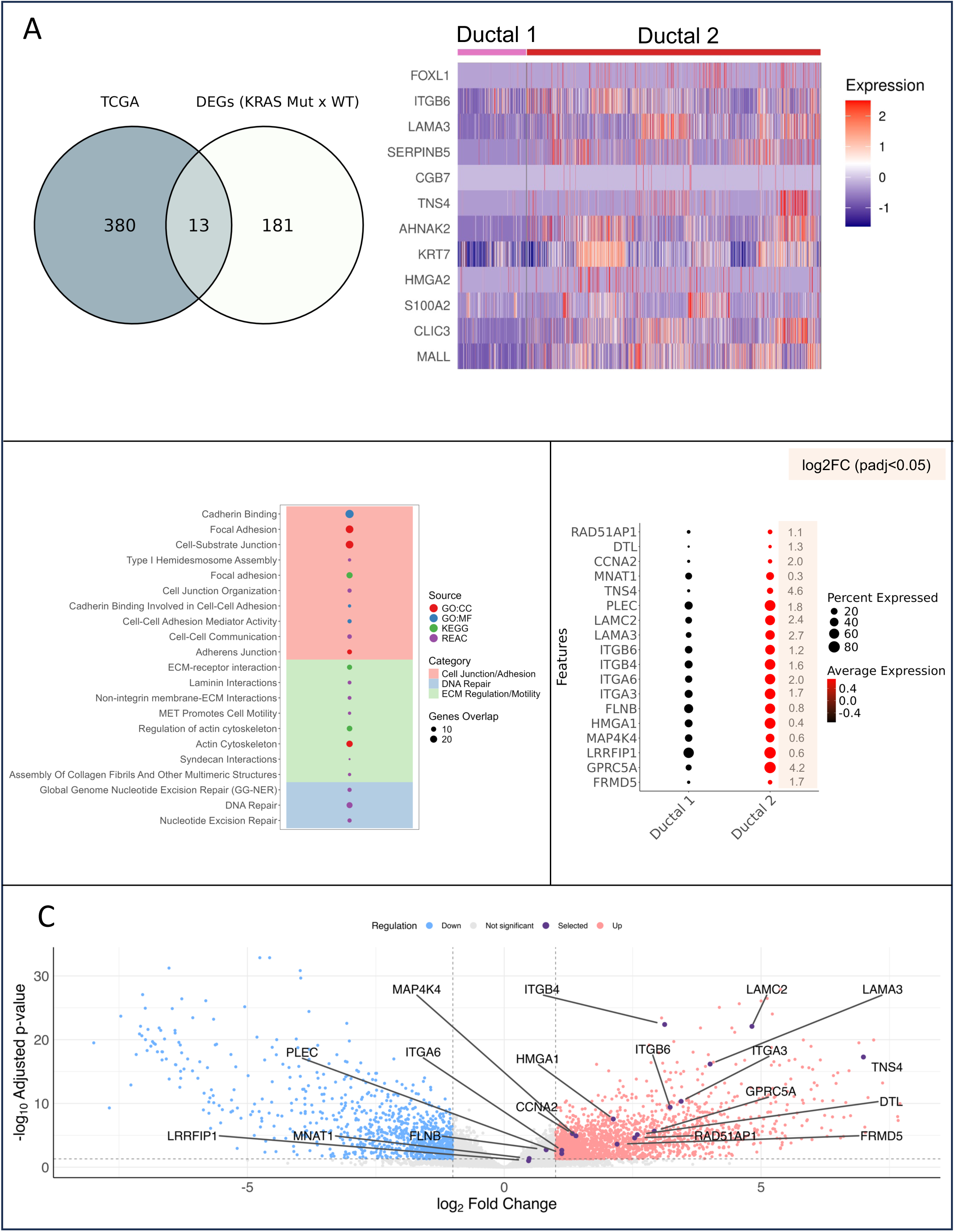
LINC00941 correlated genes are enriched in pathways related to ECM remodeling and DNA repair. (**A**) *Left:* overlap between genes significantly correlated with LINC00941 in the TCGA cohort (Spearman’s ρ ≥ 0.65) and genes differentially expressed by KRAS mutation status in the same cohort. *Right:* heatmap of the overlap genes in single-cell RNA-seq data, comparing ductal-1 (pink bar) and ductal-2 (red bar) epithelial populations. Z-score normalized expression values are shown. (**B**) Functional enrichment analysis of the top 300 LINC00941-correlated genes. Dot plot of significantly enriched pathways grouped into functional categories (Cell junction/Adhesion, ECM regulation/Motility and DNA repair, as indicated). Dot size indicates the number of LINC00941-correlated genes in each pathway; dot color denotes the functional database used for annotation. (**C**) Single-cell RNA-seq enrichment dot plot of representative genes from the pathways in (B), comparing ductal-1 and ductal-2 epithelial populations. Dot size indicates the percentage of cells in that subpopulation with detectable expression and dot color indicates the enrichment score. Differential expression was tested on normalized, log₂-transformed counts using the Wilcoxon rank-sum test, with multiple testing correction (Benjamini–Hochberg) and padj < 0.05 was considered significant. Log_2_FC values are indicated. (**D**) Volcano plot of protein-coding gene expression in paired PDAC tumors vs. adjacent pancreatic tissue (APT, n = 14). The x-axis shows log₂ fold change (PDAC vs. APT), and the y-axis shows –log₁₀ adjusted p-value. Dotted lines indicate the thresholds for significance (|log₂FC| ≥ 0.58, adjusted p < 0.05). Genes are colored by significance status (upregulated, downregulated, not significant), and LINC00941-correlated genes in enriched pathways are highlighted.

In summary, our integrative analyses establish LINC00941 as a KRAS-regulated lincRNA with functional relevance in PDAC. LINC00941 was consistently upregulated in KRAS-mutant tumors, enriched in malignant epithelial cells, and associated with poor prognosis. Functional assays demonstrated that LINC00941 promotes migration, invasion, DNA repair and chemoresistance, while transcriptomic and co-expression analyses linked it to pathways controlling DNA repair, cell motility, extracellular matrix regulation, and adhesion. These findings position LINC00941 as both a biomarker of PDAC aggressiveness and a potential therapeutic target for overcoming KRAS-driven tumor progression and treatment resistance.

## Discussion

Despite decades of research, pancreatic cancer remains a major public health challenge. It is still difficult to detect and has an extremely low survival rate, even with advancements in medicine (3). Pancreatic ductal adenocarcinoma (PDAC) is the most common form of pancreatic cancer, and KRAS mutations are found in over 90% of cases, playing a key role in oncogenic transformation (29,30). While efforts to directly inhibit KRAS have been made, and mutation-specific inhibitors for KRAS^G12C^ have generated hope, particularly in lung cancer, this approach still faces significant challenges before it can be successfully applied in clinical settings for PDAC and other cancers with different KRAS mutations (4,6). Therefore, it is essential to better understand the molecular pathways activated by oncogenic KRAS in PDAC, to identify potential new therapeutic targets.

The role of lncRNAs in the acquisition and maintenance of the malignant phenotype is becoming increasingly evident. These molecules have been implicated in all the hallmarks of cancer, and in PDAC, there is growing evidence of their significant roles (31). The interaction between lncRNAs and KRAS has been studied for several years, but findings in this area remain incomplete. Some studies describe lncRNAs that regulate the expression of this oncogene, often through miRNA sequestration (32). However, there are few reports of lncRNAs regulated by KRAS, especially in PDAC (33–39). This study contributes to filling this gap by identifying LINC00941 as a molecular target of oncogenic KRAS in PDAC, with significant clinical and functional relevance.

To identify KRAS-regulated lncRNAs, we conducted a differential expression analysis of the TCGA PDAC dataset stratified by KRAS mutation status and identified 49 differentially expressed intergenic lncRNAs. Although few KRAS-regulated lncRNAs in PDAC have been characterized, our analysis identified *UCA1*, a known oncogenic lncRNA previously shown to be upregulated by KRAS in PDAC (36,37). Additionally, we found that the expression of known PDAC-associated oncogenic lncRNAs, such as LINC01559 (40), LINC01133 (41), LINC00941 (42) and LINC00462 (43) was significantly associated with KRAS status. These findings suggest that these lncRNAs are likely downstream targets of KRAS in PDAC.

In contrast, the KRAS-associated lncRNAs identified in this study were not differentially expressed according to KRAS mutation status in a TCGA cohort of colorectal cancer patients (33). This observation underscores the context-dependent nature of lncRNA expression and function and supports the notion that the lncRNAs identified here may act as PDAC-specific KRAS biomarkers.

To further investigate whether KRAS-associated lncRNA expression correlates with specific KRAS mutations, we examined the expression of the ten most highly expressed KRAS-associated lncRNAs in the TCGA PDAC cohort stratified by KRAS mutation type. This analysis is relevant since specific KRAS mutations are known to differently affect downstream signaling pathways and therapeutic response. For instance, PDAC KRAS^G12D^ cells are 10 times more sensitive to MEK inhibition than PDAC KRAS^G12C^ cells (44). However, our results showed that lncRNA upregulation in KRAS-mutant tumors did not depend on specific KRAS mutations. This suggests that these lncRNAs may serve as general biomarkers for KRAS-mutant PDAC, independent of mutation subtype. Furthermore, if therapeutically actionable, they could be targeted alongside other therapies to improve outcomes, particularly in cases involving KRAS mutations with poor therapeutic response.

Among the ten KRAS-associated lncRNAs initially selected, LINC00941 was positively regulated by KRAS in both gain- and loss-of-function models in pancreatic cells, underscoring its potential as a KRAS effector. LINC00941 has been implicated in various cancers, including PDAC (45,46). In a previous study, we identified LINC00941 as upregulated in PDAC (14). Here, we confirmed that LINC00941 expression is associated with poor survival in the TCGA PDAC dataset and in the ICGC cohort and demonstrated its enrichment in the malignant epithelial cell population, indicating a cancer cell-intrinsic function. Functional analyses showed that LINC00941 silencing reduces PDAC cell migration and invasion, two key traits of malignant behavior, while having no evident effect on cell viability or death. These findings are supported by other studies showing that LINC00941 is upregulated in PDAC, is associated with worse prognosis and regulates PDAC cell malignant traits (47–50).

We had previously identified that LINC00941 was co-expressed with DNA repair genes in PDAC (14). We now show that LINC00941 functionally promotes DNA repair and upregulates core DNA repair genes. Among these, the most consistently regulated genes were PPP4C, a phosphatase that facilitates homologous recombination via PLK1 dephosphorylation (51), and BACH1, a BRCA1-interacting helicase involved in double-strand break repair (52,53). Notably, both are significantly enriched in malignant type 2 ductal cells. BACH1 expression is also increased in KRAS mutant patients and PPP4C is associated with cancer progression and poor prognosis in other tumor types (54).

Enhanced DNA repair is a known molecular mechanism that contributes to chemoresistance (25). Consistently, we found that LINC00941 also promotes gemcitabine chemoresistance. This finding is supported by reports showing that other noncoding RNAs were found to enhance gemcitabine resistance through activation of DNA repair pathways (55,56).

Beyond DNA repair, our findings suggest that LINC00941 contributes to PDAC chemoresistance through regulation of downstream effectors involved in both drug efflux and apoptotic resistance. LINC00941 knockdown reduced the expression of ABCG2, an ATP-binding cassette transporter that mediates efflux of several molecules, including anticancer drugs (57) and is implicated in multidrug resistance and tumor progression (57), as well as BIRC5 (Survivin), a member of the Inhibitor of Apoptosis (IAP) family (58). Although both genes showed expression trends consistent with upregulation in PDAC bulk tumors and in malignant PDAC cells, only BIRC5 was significantly elevated and strongly associated with reduced overall survival in PDAC patients.

The relevance of BIRC5 is supported by previous studies demonstrating its role in promoting resistance to gemcitabine in PDAC models (28,59,60). In addition, BIRC5 is markedly overexpressed in many tumors, including PDAC, with high levels correlating with aggressive tumour features(61). Mechanistically, BIRC5 inhibits caspase-dependent apoptosis and facilitates cell cycle progression, thereby allowing cancer cells to evade chemotherapy-induced cell death (58). Interestingly, BIRC5 has also been linked to enhanced DNA damage repair and mitotic regulation, further reinforcing its role in sustaining tumor cell survival under genotoxic stress (62,63). High BIRC5 expression has been consistently reported across solid tumors as a marker of aggressive disease and poor prognosis (58), which is in agreement with our survival analyses in PDAC. Thus, in our results, the stronger upregulation of BIRC5 in comparison to ABCG2, its significant prognostic association, and its known capacity to suppress apoptosis make it likely that LINC00941’s effect on chemoresistance is mediated in large part through apoptosis inhibition, in addition to drug efflux. LINC00941-mediated upregulation of ABCG2 and BIRC5 correlates with the aggressive and chemoresistant phenotype frequently observed in PDAC tumors.

In addition, the set of enriched pathways identified from the top LINC00941-correlated genes supports a model in which LINC00941 may promote PDAC aggressiveness and chemoresistance via two major axes: (1) enhancing invasive/motile behavior through ECM and adhesion dynamics, and (2) increasing the ability to repair genotoxic damage and survive DNA-targeting therapies. Among the genes driving these enrichments, ITGA3 and RAD51AP1 stand out as both upregulated in PDAC and significantly associated with patient survival. ITGA3, an integrin alpha subunit involved in cell–ECM adhesion and signaling, has been linked to invasion, metastasis, and poor prognosis, with functional studies showing that its inhibition reduces PDAC growth and invasiveness (64,65). RAD51AP1, a RAD51-interacting protein that promotes homologous recombination, is likewise upregulated in PDAC and associated with poor survival, likely by enhancing DNA repair capacity and conferring resistance to genotoxic therapies (66,67). Together, these genes exemplify how LINC00941-associated pathways converge on both ECM remodeling and DNA repair, reinforcing its connection to aggressive, therapy-resistant PDAC.

In conclusion, our study identifies LINC00941 as a key KRAS-regulated lncRNA that drives PDAC aggressiveness by enhancing migration, invasion, DNA repair, and chemoresistance. Mechanistically, we show that LINC00941 upregulates core DNA repair factors such as PPP4C and BACH1, and chemoresistance mediators including ABCG2 and BIRC5. Among these, BIRC5 stands out as a strong predictor of poor survival and a likely major effector of LINC00941-mediated resistance to gemcitabine. The integration of these validated targets with pathway analyses strengthens the model that LINC00941 promotes tumor progression through both apoptosis inhibition and DNA repair enhancement. These findings highlight LINC00941 as a clinically relevant downstream effector of KRAS signaling in PDAC and suggest that therapeutic strategies aimed at inhibiting LINC00941 or its key effectors may offer dual benefits of sensitizing KRAS-induced PDAC tumors to chemotherapy and limiting metastatic progression.

## Ethics approval and consent to participate

The study involving human samples was approved by the Ethics Committee of the AC Camargo Cancer Center (CAAE: 15059213.0.0000.5432). All procedures were perfomed in accordance with the 1964 Helsinki declaration and its later amendments or comparable ethical standards. Informed consent was obtained from all participants included in the study.

## Author contributions

T.B.C. participated in the study design, performed qPCR validation of lincRNAs in cell lines and patients, performed all functional studies of LINC00941, analyzed results and drafted the manuscript. G.L.F. and R.B.M. conducted all bioinformatics analyses under E.M.R.’s and D.S.B’s supervision. L.A.H. generated and performed western blotting of HPDE/HPDE-KR cells, S.M.G-F performed western blottings of AsPC-1 cells. Y.M. supervised by F.L.F helped with comet assay experiments and analysis. L.R. supervised by N.C. helped with cell death experiments, image acquisition and analysis. E.L. participated in the study design and critically revised the manuscript. E.M.R. participated in the study design, supervised bioinformatics analysis and acted as a co-advisor to T.B.C. and G.L.F. D.S.B. conceived the study, participated in its design and coordination, and helped to draft and revise the manuscript. All authors have read and agreed to the published version of the manuscript.

## Data availability statement

The datasets generated during and/or analysed during the current study are available from the corresponding author on reasonable request.

