## Supplementary Figure legends for "LncRNA LINC00941 Links Oncogenic KRAS Signaling to Aggressiveness and Chemoresistance in Pancreatic Cancer"

**Supplementary Figure 1. Expression of selected lincRNAs according to *KRAS* mutation status in the ICGC PDAC cohort.** Dot plots comparing $log_2$-transformed CPM-normalized counts of the indicated lincRNAs in $KRAS^{MUT}$ (n = 9) and $KRAS^{WT}$ (n = 89) tumors in the ICGC PDAC cohort (PACA-AU, n = 98). Statistical significance was assessed using two-tailed Wilcoxon rank-sum tests, with $p < 0.05$ considered significant.

**Supplementary Figure 2. LINC00941 depletion affects the expression of LINC00941-correlated DNA repair genes in PANC-1 cells.** PANC-1 cells were transfected with non-targeting control siRNA (siCTRL) or a SMARTpool siRNA targeting LINC00941 (si941). **(A)** qRT-PCR analysis of LINC00941 and selected DNA-repair genes 48 h after transfection (n = 3 independent experiments). Data are shown as fold-change relative to siCTRL (mean ± SD). HMBS was used as endogenous control. Bars colored red indicate genes positively correlated with LINC00941; blue indicates negatively correlated genes. Statistical significance was assessed using paired one-tailed (LINC00941) or two-tailed (DNA repair genes) Student's *t*-test. Significance levels are denoted as *$p < 0.05$, **$p < 0.01$. Exact *p*-values and Cohen's *d* effect sizes are indicated.

**Supplementary Figure 3. High expression of LINC00941-correlated genes in the TCGA cohort ITGA3 and RAD51AP1 is associated with poor survival in PDAC.** Kaplan–Meier survival analysis based on ITGA3 (left) and RAD51AP1 (right) expression in the TCGA (PAAD-US, n = 145) and ICGC (PACA-AU, n = 87) cohorts as indicated. Patients were stratified into high- and low-expression groups using the median expression cutoff. Differences in survival were evaluated using the log-rank test; log-rank *p*-values and hazard ratios (HR) with 95% confidence intervals are shown.
