## Supplementary material for "LncRNA LINC00941 Links Oncogenic KRAS Signaling to Aggressiveness and Chemoresistance in Pancreatic Cancer": Suplementary Table 1

**SUPPLEMENTARY TABLE 1:** Sequence of primers used for qRT-PCR

| Gene | Forward sequence (5'-3') | Reverse sequence (5'-3') |
|---|---|---|
| LINC00941 | TTTTGTGTCCAAGCCCCAGA | TGGGAGACTGAGATGGAAGG |
| AC006262.5 | TCCTCAGCCCAGCCGAAG | AGAGGGTGTTCACATTGCTTT |
| KRAS | CCCAGGTGCGGGAGAGA | CAGCTCCAACTACCACAAGTTT |
| BACH1 | GAACCCCAGCAAGAACCTTG | TCCGTTGTGCATTGAATGGC |
| TERF2IP | CCTCCTCCCAGAAGCTCAAG | CAGGCAAATCTGGAGTTCTCT |
| CDK7 | GCACACCAACTGAGGAACA | TCTCCTGCTGCACTGAAGAT |
| ATRX | AGCATTAAGTAGACAAGCCAGC | CTGATTGTACTGCTGCTGGA |
| TOP2A | TGGTGGCAAGGATTCTGCTA | CATTCAGGCTCAACACGCTG |
| BIRC5 | GGACCACCGCATCTCTACAT | GAAACACTGGGCCAAGTCTG |
| SPIDR | AGAATGCCCATCCTTTCCAG | AGGTGCTTTTCTGTAATCGTC |
| PPP4C | ATGACCTCAAAGAGCTGTTCAG | TAGAAGCCACGGTCCACAAAG |
| GAPDH | GAGCCGCATCTTCTTTTGC | CCATGGTGTCTGAGCGATGT |
| HMBS | GGCAATGCGGCTGCAA | GGGTACCCACGCGAATCAC |
