## Supplementary figures and images for "LncRNA LINC00941 Links Oncogenic KRAS Signaling to Aggressiveness and Chemoresistance in Pancreatic Cancer"

### Supplementary Figure 1

#### SUPPLEMENTARY FIGURE 1

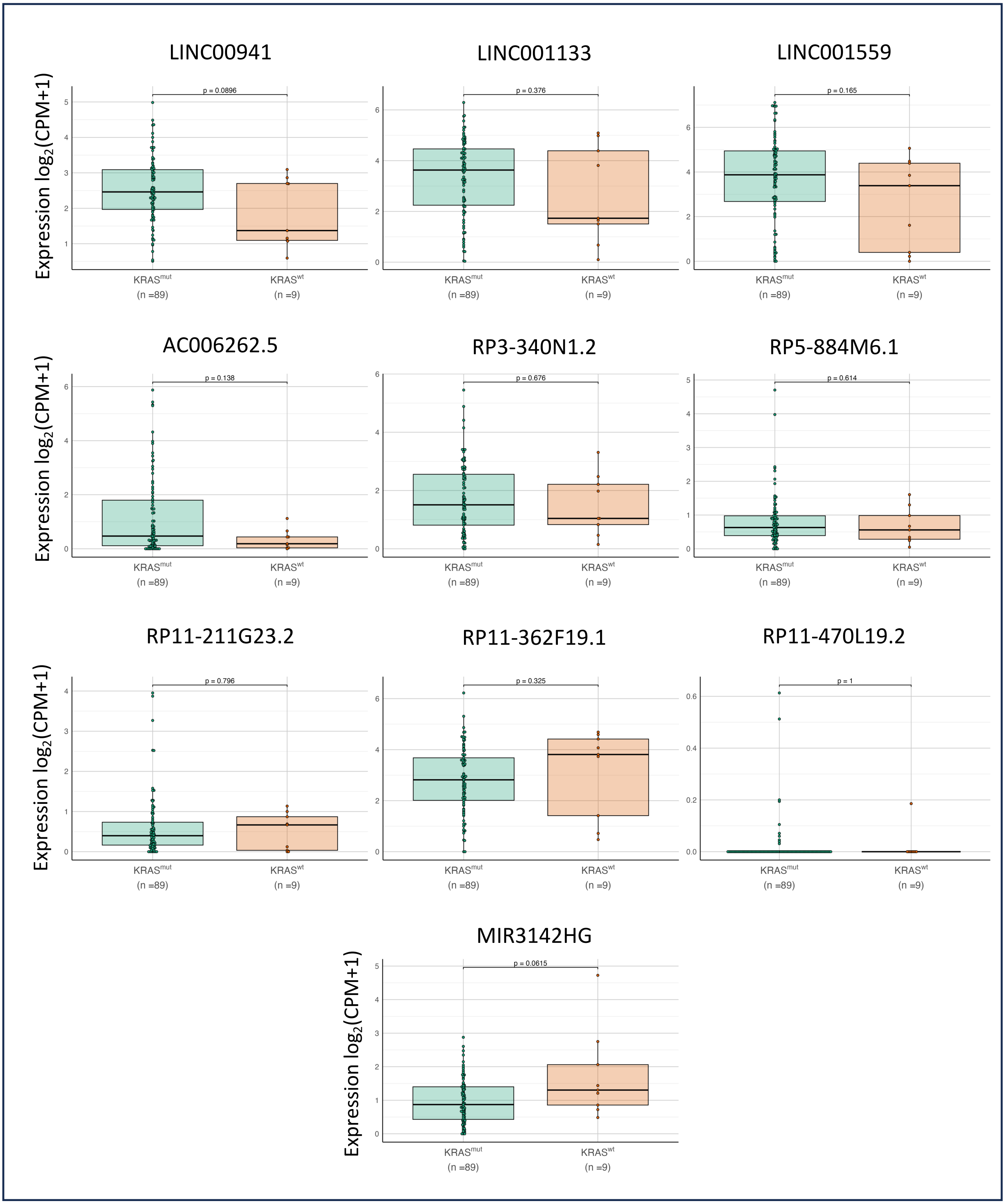

### Supplementary Figure 2

#### SUPPLEMENTARY FIGURE 2

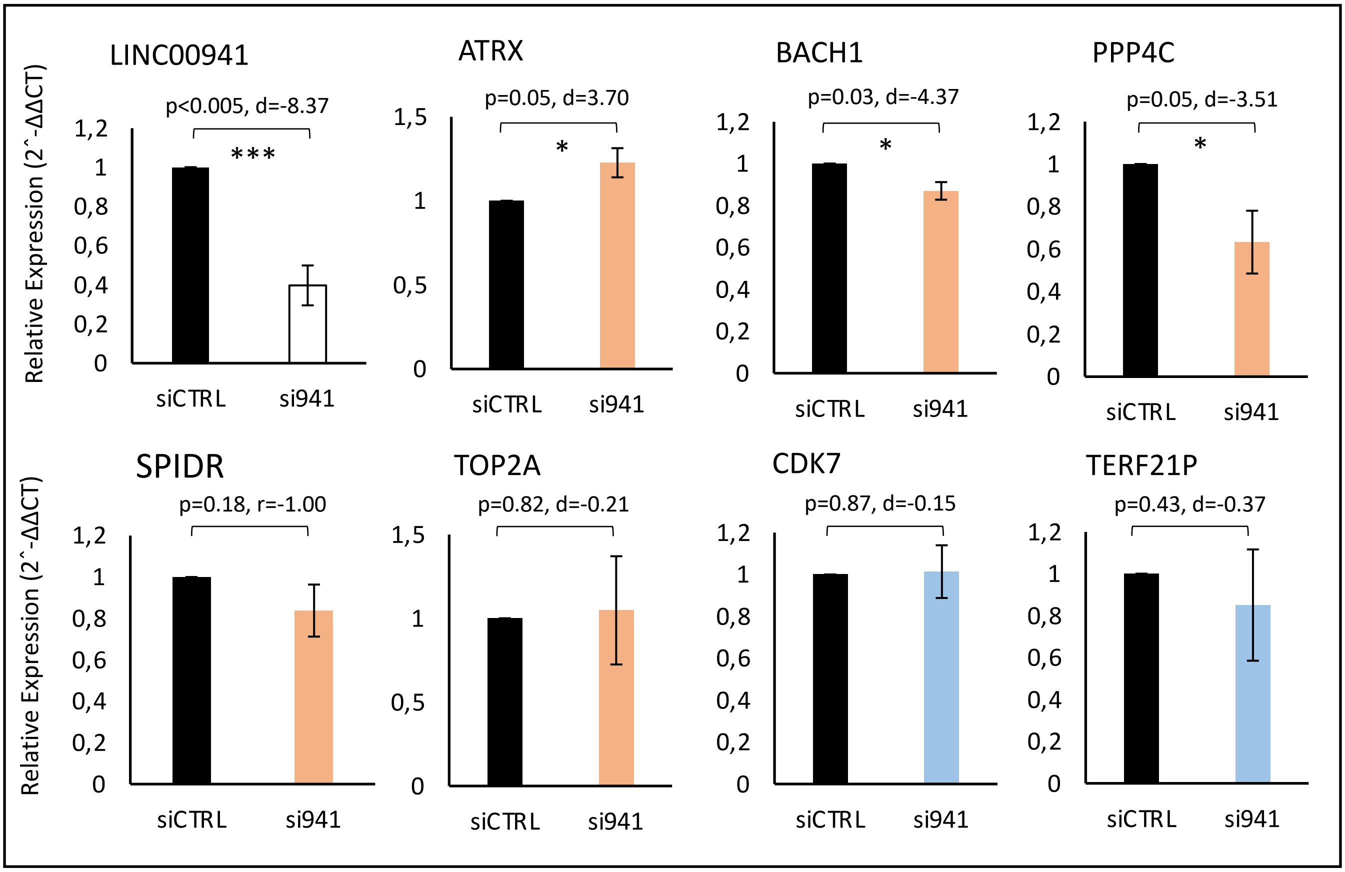

### Supplementary Figure 3

#### SUPPLEMENTARY FIGURE 3

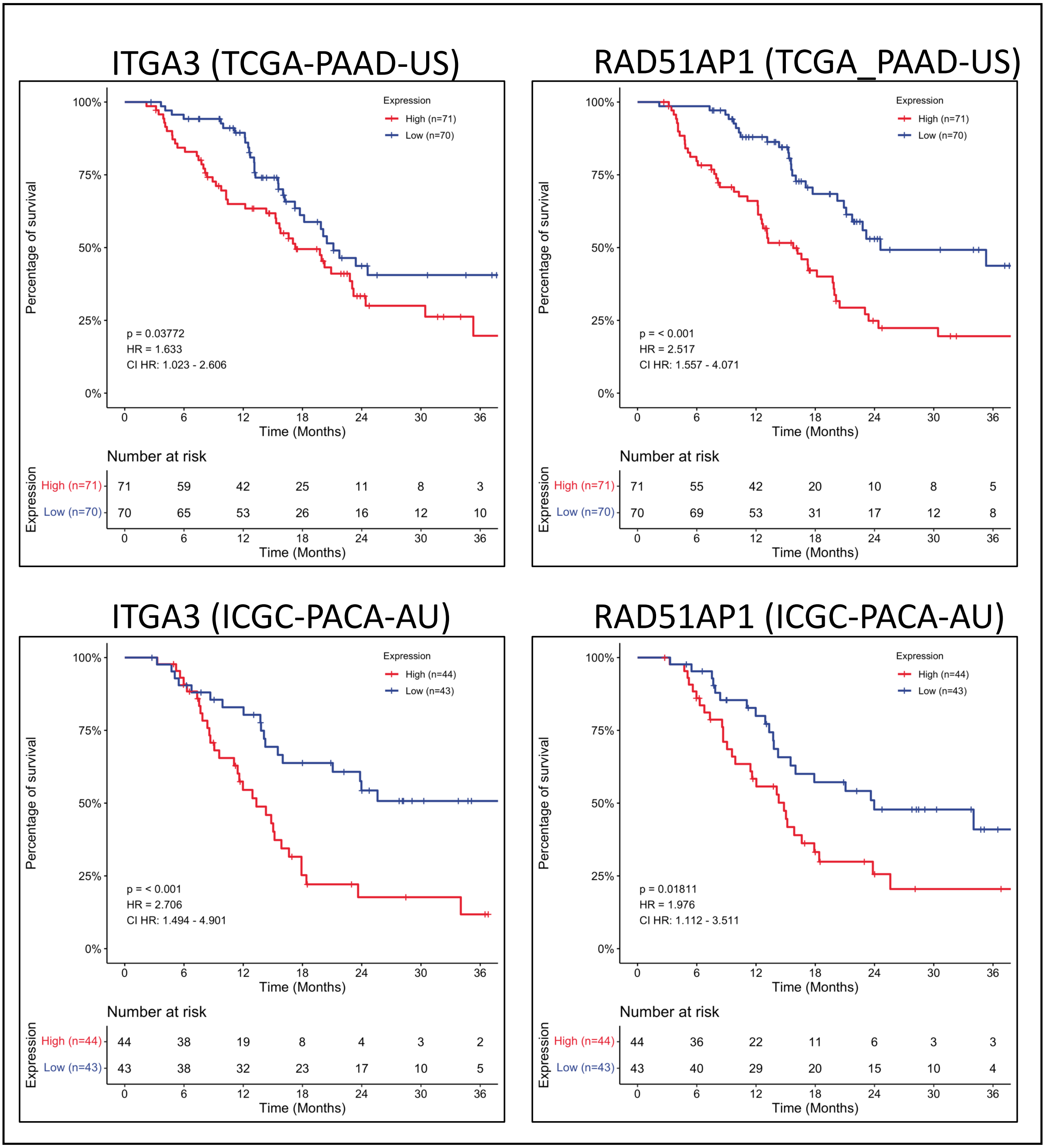
